# Fusarium Protein Toolkit: AI-powered tools to combat fungal threats to agriculture

**DOI:** 10.1101/2024.04.30.591916

**Authors:** Hye-Seon Kim, Olivia C. Haley, John L. Portwood, Stephen Harding, Robert H. Proctor, Margaret R. Woodhouse, Taner Z. Sen, Carson M. Andorf

## Abstract

**Background:** The fungal genus *Fusarium* poses significant threats to food security and safety worldwide because it consists of numerous species that cause destructive diseases in crops, as well as mycotoxin contamination. The adverse effects of climate change are exacerbating some existing threats and causing new problems. These challenges highlight the need for innovative solutions, including the development of advanced tools to identify targets to control crop diseases and mycotoxin contamination incited by *Fusarium*.

**Description:** In response to these challenges, we developed the Fusarium Protein Toolkit (FPT, https://fusarium.maizegdb.org/), a web-based tool that allows users to interrogate the structural and variant landscape within the *Fusarium* pan-genome. FPT offers a comprehensive approach to understanding and mitigating the detrimental effects of *Fusarium* on agriculture. The tool displays both AlphaFold and ESMFold-generated protein structure models from six *Fusarium* species. The structures are accessible through a user-friendly web portal and facilitate comparative analysis, functional annotation inference, and identification of related protein structures. Using a protein language model, FPT predicts the impact of over 270 million coding variants in two of the most agriculturally important species, *Fusarium graminearum*, which causes Fusarium head blight and trichothecene mycotoxin contamination of cereals, and *F. verticillioides*, which causes ear rot and fumonisin mycotoxin contamination of maize. To facilitate the assessment of naturally occurring genetic variation, FPT provides variant effect scores for proteins in a *Fusarium* pan-genome constructed from 22 diverse species. The scores indicate potential functional consequences of amino acid substitutions and are displayed as intuitive heatmaps using the PanEffect framework.

**Conclusion:** FPT fills a knowledge gap by providing previously unavailable tools to assess structural and missense variation in proteins produced by *Fusarium*, the most agriculturally important group of mycotoxin-producing plant pathogens. FPT will deepen our understanding of pathogenic mechanisms in *Fusarium*, and aid the identification of genetic targets that can be used to develop control strategies that reduce crop diseases and mycotoxin contamination. Such targets are vital to solving the agricultural problems incited by *Fusarium*, particularly evolving threats affected by climate change. By providing a novel approach to interrogate *Fusarium*-induced crop diseases, FPT is a crucial step toward safeguarding food security and safety worldwide.

## 1 Background

Understanding the dynamics of the interactions of *Fusarium* species and their crop hosts on a molecular level is critical for developing effective control strategies to mitigate the adverse economic and health impacts caused by these fungi. Fusarium head blight (FHB) of wheat and barley, which is caused primarily by *Fusarium graminearum*, leads to significant yield losses and mycotoxin contamination, with economic losses in the U.S. estimated at $1.5 billion between 2015 and 2016 [1]. Similarly, Fusarium ear rot of maize, caused by *Fusarium verticillioides*, not only reduces yield and quality but also results in mycotoxin contamination, which together adversely affects corn producers and processors. Climate change is expected to increase the susceptibility of crops to *Fusarium*-incited diseases and mycotoxin contamination, thereby amplifying their negative economic impact [2]. This is because climate change alters temperature and precipitation patterns, creating favorable conditions for *Fusarium* growth and reproduction, and increasing the likelihood of pathogen-crop interactions. Additionally, climate-related stress can weaken crop immune systems [3], making them more vulnerable to *Fusarium* infection. An integrated approach that leverages scientific knowledge and technological innovation is essential to address the negative impacts of *Fusarium* on agriculture.

A significant gap in knowledge of *Fusarium* is the absence of a comprehensive resource for searching and visualizing protein structures and exploring the functional consequences of amino acid substitutions within these proteins. This limitation hinders our ability to understand the molecular mechanisms underlying Fusarium pathogenesis and toxin production, making it more challenging to develop effective strategies for battling Fusarium-related diseases. Recent advancements, notably AlphaFold [4] and other technologies, have made it possible to predict accurate protein structures for most amino acid sequences, broadening the possibilities for comparative structural studies. The AlphaFold Protein Structure Database [5] now offers protein structures for over 200 million proteins, covering the whole proteomes of many species. Additionally, new tools like FoldSeek[6], which is based on a novel structural alphabet and tertiary interactions, offer faster protein alignment capabilities than previous methods.

Extending these tools to predict functional effects of missense mutations—i.e., mutations in codons that cause an amino acid substitution in the corresponding protein—is crucial for understanding the molecular mechanisms underlying Fusarium evolution, pathogenesis, and adaptation, and could potentially reveal new targets for disease management and crop improvement. Missense mutation approaches have been explored in research on human biology [7–9] but not *Fusarium* biology. The AlphaMissense [7] resource leverages structural and evolutionary information to classify all possible missense mutations in the human genome as either benign or likely pathogenic (i.e., deleterious). The GEMME [8] and the esm-variants [9] workflows are other examples where alignment-based strategies and large-scale protein language models predict mutational outcomes across numerous protein families and the entire human proteome, respectively. However, a tool that facilitates exploration of the diversity of publicly available Fusarium proteomes and predicts potential functional consequences of missense changes in codons is yet to be developed.

A new resource tailored for *Fusarium* research should incorporate innovative technologies to enable comprehensive exploration of *Fusarium* protein structures and missense variants. Such a resource would not only facilitate the identification and visualization of protein structures but also enable detailed analysis of the functional implications of genetic variants. By combining structural predictions with variant effect prediction technologies, this tool would greatly enhance our understanding of *Fusarium* pathogenicity and potential targets for control. Additionally, it should provide an intuitive platform for comparative genomics, enabling researchers to compare *Fusarium* strains and identify key genetic targets for developing effective disease management strategies. Ultimately, this resource would represent a significant advancement in *Fusarium* research, offering insights into the genetic basis of variation in these pathogens and informing the development of innovative control measures.

Here, we introduce a new database resource called the Fusarium Protein Toolkit (FPT - https://fusarium.maizegdb.org/) which offers a suite of tools for exploring protein structures, variant effects, and annotated effector proteins from the genus *Fusarium*. FPT leverages the frameworks and tools developed for maize (*Zea mays*) by the Maize Genetics and Genomics Database (MaizeGDB) [10, 11]. This shared infrastructure sets a foundation for future studies focused on host-pathogen interactions and can be effortlessly implemented in other biological databases such as GrainGenes [12]. FPT can be used as a stand-alone resource to explore *Fusarium* proteins or to facilitate comparisons of protein structures and functionalities between *Fusarium* and maize. These comparisons can enable researchers to gain an in-depth understanding of the molecular interactions between these organisms. Such insights can help inform the development of targeted strategies to control crop diseases and mycotoxin contamination.

## 2 Construction and content

### Data Sources and Curation

The main features of the Fusarium Protein Toolkit (FPT) include predicted three-dimensional (3D) structures of proteins, which aid determination of function, and predictions of whether amino acid substitutions in orthologous proteins impact protein function. For the development of FPT, the following primary data types were collected or generated, and curated for *Fusarium* proteomes: 1) *E*ffector proteins, 2) protein structure models, 3) pan-genome framework with protein alignments, orthology groups, and variant effect scores. An overview of the 22 *Fusarium* species integrated into the FPT can be found in Table 1 and a list of the data and tools available for each species is found in Table 2.

**Table 1.** Overview of *Fusarium* species in the Fusarium Protein Toolkit.

| Full name | Taxonomy | UniProt | Protein Count |
| --- | --- | --- | --- |
| *Fusarium acutatum* | 78861 | UP000536711 | 14,072 |
| *Fusarium avenaceum* | 40199 | UP000782241 | 11,232 |
| *Fusarium coffeatum* | 231269 | UP000253153 | 11,778 |
| *Fusarium culmorum* | 5516 | UP000241587 | 12,350 |
| *Fusarium duplospermum* | 1325734 | UP000288168 | 16,262 |
| *Fusarium flagelliforme* | 2675880 | UP000265631 | 13,039 |
| *Fusarium fujikuroi* | 1279085 | UP000016800 | 14,792 |
| *Fusarium graminearum* | 229533 | UP000070720 | 16,422 |
| *Fusarium mangiferae* | 192010 | UP000184255 | 15,798 |
| *Fusarium musae* | 1042133 | UP000827133 | 13,672 |
| *Fusarium odoratissimum* | 1089451 | UP000030685 | 19,807 |
| *Fusarium oxysporum* | 426428 | UP000009097 | 23,111 |
| *Fusarium poae* | 36050 | UP000091967 | 14,048 |
| *Fusarium proliferatum* | 1227346 | UP000183971 | 16,122 |
| *Fusarium pseudograminearum* | 1028729 | UP000007978 | 12,448 |
| *Fusarium redolens* | 48865 | UP000720189 | 17,005 |
| *Fusarium vanettenii* * | 660122 | UP000005206 | 15,711 |
| *Fusarium sporotrichioides* | 5514 | UP000266152 | 11,960 |
| *Fusarium subglutinans* | 42677 | UP000547976 | 14,039 |
| *Fusarium tjaetaba* | 1567544 | UP000530670 | 14,180 |
| *Fusarium venenatum* | 56646 | UP000245910 | 13,945 |
| *Fusarium verticillioides* | 334819 | UP000009096 | 17,876 |

This table provides a list of the 22 *Fusarium* species accessible via the Fusarium Protein Toolkit. It includes details for each species, such as the full species name (from UniProt), the NCBI Taxonomy Database ID, the UniProt Proteome ID, and the total number of proteins in each proteome from UniProt.
\*Formerly *F. solani* f. sp. *lisi* and *Nectria haematococca*.

**Table 2.** Data Availability for *Fusarium* species in the Fusarium Protein Toolkit.

| **Name** | **FASTA** | **PDB** | **Variant scores** | **Effectors** | **FoldSeek** | **PanEffect** |
| --- | --- | --- | --- | --- | --- | --- |
| *F. acutatum* | Y | N | Y | N | N | Y |
| *F. avenaceum* | Y | N | Y | N | N | Y |
| *F. coffeatum* | Y | N | Y | N | N | Y |
| *F. culmorum* | Y | N | Y | N | N | Y |
| *F. duplospermum* | Y | N | Y | N | N | Y |
| *F. flagelliforme* | Y | N | Y | N | N | Y |
| *F. fujikuroi* | Y | Y | Y | Y | Y | Y |
| *F. graminearum* | Y | Y | Y | Y | Y | Y |
| *F. mangiferae* | Y | N | Y | N | N | Y |
| *F. musae* | Y | N | Y | N | N | Y |
| *F. odoratissimum* | Y | N | Y | N | N | Y |
| *F. oxysporum* | Y | Y | Y | Y | Y | Y |
| *F. poae* | Y | N | Y | N | N | Y |
| *F. proliferatum* | Y | Y | Y | Y | Y | Y |
| *F. pseudograminearum* | Y | N | Y | N | N | Y |
| *F. redolens* | Y | N | Y | N | N | Y |
| *F. vanettenii\** | Y | Y | Y | Y | Y | Y |
| *F. sporotrichioides* | Y | N | Y | N | N | Y |
| *F. subglutinans* | Y | N | Y | N | N | Y |
| *F. tjaetaba* | Y | N | Y | N | N | Y |
| *F. venenatum* | Y | N | Y | N | N | Y |
| *F. verticillioides* | Y | Y | Y | Y | Y | Y |

This table outlines the datasets available for download for the 22 *Fusarium* species featured in the Fusarium Protein Toolkit. It details the types of data accessible (including FASTA protein sequences, ESMFold protein structures in the PDB format, and variant effect scores) and specifies which species have been integrated into the Effector tables, FoldSeek search, and PanEffect tools. \*Formerly *F. solani* f. *sp. pisi* and *Nectria haematococca*.

### Prediction of effector proteins

Like other plant pathogenic fungi, *Fusarium* species secrete small proteins known as effectors that confer the ability to overcome plant defenses and, thereby, enable the fungi to cause disease. Genomic data from fungal plant pathogens has been used to identify effectors and determine the underlying mechanism by which they impact plant defenses. However, genomic bases studies of *Fusarium* are limited and there are no genome-wide analyses of effectors from multiple species. To address this knowledge gap, we used a combination of genome sequence analyses and machine learning approaches to develop a five-step computational pipeline to identify effectors, assess ortholog distribution, and predict potential function. The genome sequences for six representative *Fusarium* species (*F. fujikuroi, F. graminearum, F. oxysporum, F. proliferatum, F. vanettenii*, and *F. verticillioides*) were downloaded from the NCBI Reference Sequence Database (RefSeq) [13] An overview of numbers of predicted proteins, effector proteins, and secreted proteins in these species is provided in Table 3.

**Table 3.** Overview of the *Fusarium* species used in the effector annotation pipeline.

| **Species** | **NCBI RefSeq** | **Total proteins** | **EffectorP proteins** | **SecretSanta proteins** |
| --- | --- | --- | --- | --- |
| *F. fujikuroi* | GCF_900079805.1 | 14,813 | 4,391 | 390 |
| *F. graminearum* | GCF_000240135.3 | 13,313 | 4,424 | 290 |
| *F. oxysporum* | GCF_000149955.1 | 20,925 | 6,799 | 462 |
| *F. proliferatum* | GCF_900067095.1 | 16,125 | 4,711 | 439 |
| *F. vanettenii* * | GCF_000151355.1 | 15,708 | 4,002 | 308 |
| *F. verticillioides* | GCA_000149555.1 | 15,869 | 4,884 | 412 |

The table shows the six *Fusarium* species used to calculate putative effector proteins along with the NCBI RefSeq ID, the total number of proteins downloaded from RefSeq, the number of proteins annotated as likely effectors by EffectorP, and the number of effector proteins identified as likely secretors by SecretSanta. The proteins in the final column are used as the curated dataset in the Fusarium Protein Toolkit website. \*Formerly *F. solani* f. sp. *pisi* and *Nectria haematococca*.

The five-step effector protein prediction pipeline used on the six representative *Fusarium* species:

- Step 1: The effector prediction program EffectorP (version 2.0 and 3.0) [14, 15] was used to predict which proteins were effectors. This program identified 4,002 to 6,799 effector proteins in each of the six species (Table 3).
- Step 2: The secreted protein *in silico* identification program SecretSanta [16] is an efficient secretome prediction workflow that combines multiple individual command line and/or web-interfaces tools such as SignalP [17], TargetP [18], TMHMM [19], TOPCONS [20] for signal peptide, motifs, and transmembrane domains prediction and WolfPsort [21] for the protein subcellular localization prediction. SecretSanta was used to identify candidate effectors that are likely to be secreted from the fungus into the plant cell. The size of the predicted secretomes ranged from 308 to 463 proteins (Table 3).
- Step 3: The ortholog identification program OrthoFinder [22] (with default parameters) was used to identify and analyze the distribution of putative secreted effectors for the six *Fusarium* species.
- Step 4: The localization identification program LOCALIZER [23] identified potential subcellular localization signals on the putative secreted effectors that could direct them to the nucleus, chloroplast, or mitochondria.
- Step 5: The functional annotation program OmicsBox [24] predicted the functions of the putative effectors. OmicsBox utilizes CloudBlast software, which runs a Basic Local Alignment Search Tool (BLAST) analysis [25] against the NCBI-NR protein database. OmicsBox was used to predict the putative function of effector proteins by generating *Fusarium* secretome profiles for the six representative species and utilizing them as the primary, curated dataset for FPT. This dataset includes integrated profile list tables that provide detailed information and links to the FoldSeek and PanEffect tools for further analysis.

### Protein structure models

Protein structure models for the six representative species (*F. graminearum, F. fujikuroi, F. oxysporum, F. proliferatum, F. vanettenii*, and *F. verticillioides*) and four outgroups (*Arabidopsis thaliana, Homo sapiens, Saccharomyces cerevisiae, Schizosaccharomyces pombe*) were downloaded from the AlphaFold Protein Structure Database (https://alphafold.ebi.ac.uk/, Release 4). Additionally, ESMFold [26] was used to generate an independent set of 3D models for each proteome. For individual proteins, the 3D models derived from the two programs can differ because AlphaFold uses a deep learning approach that relies on multiple-sequence alignments, while ESMFold uses a protein language model. The use of the two independent sets of 3D models is expected to enhance FPT’s structural analysis capabilities. Figure 1 provides a comparison of the average per residue confidence scores for the AlphaFold and ESMFold models for each of the six *Fusarium* proteomes. The software FoldSeek[6] facilitated the generation of both structural and sequence alignments. With FoldSeek, the 15,911 and 17,356 predicted protein structures for two agriculturally important species *F. graminearum* and *F. verticillioides*, respectively, were aligned to the predicted protein structures from the other four *Fusarium* species and four outgroup species. The resulting output was modified to highlight the top 25 hits (based on the highest alignment score) from each species, and to incorporate functional annotations from UniProt, species information, and a visual blue/red color gradient for intuitive interpretation of results [10, 27].

**Figure 1.**
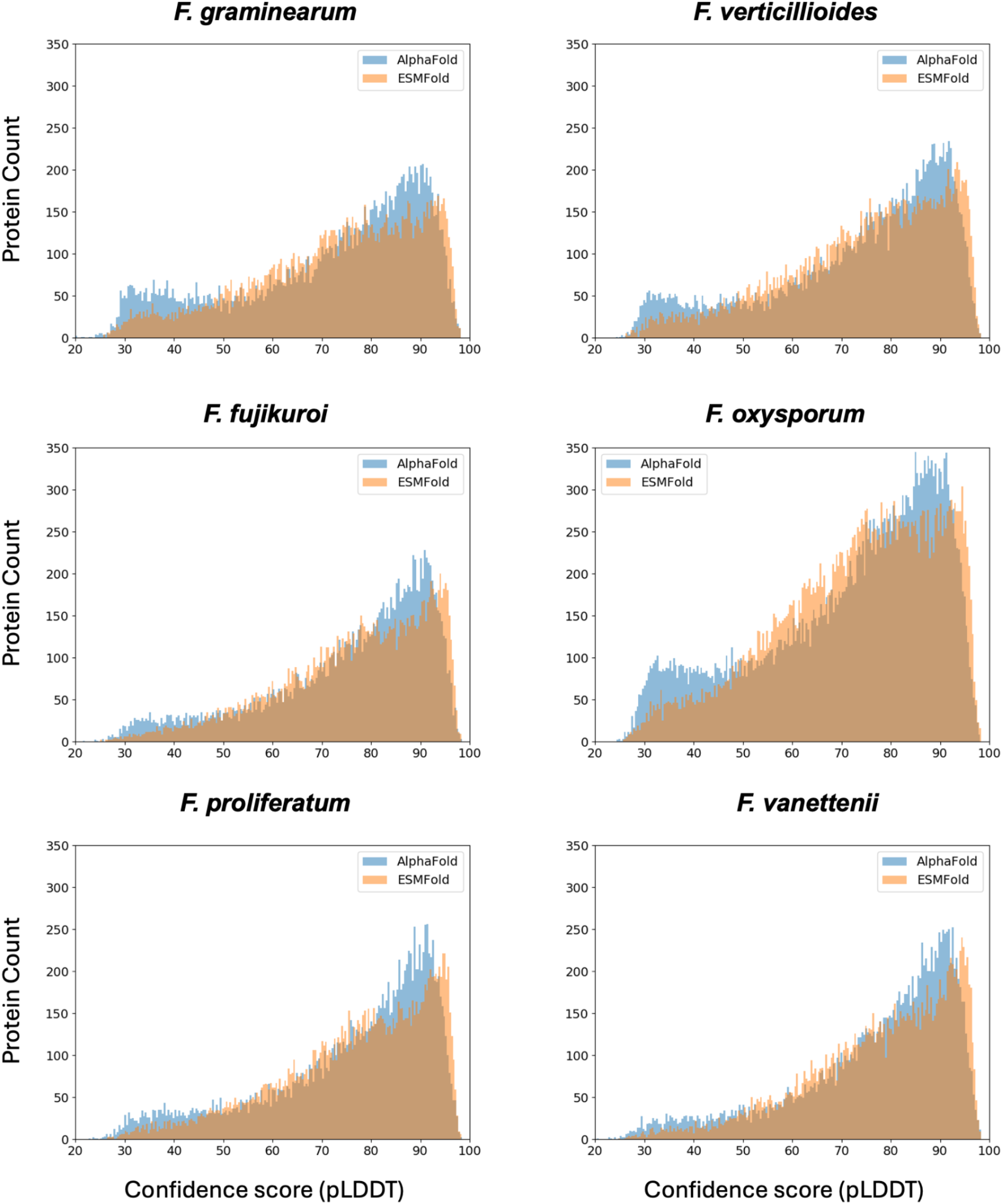
Comparison of the distribution of confidence scores from AlphaFold and ESMFold. The Fusarium Protein Toolkit provides both AlphaFold and ESMFold structures. AlphaFold is a deep learning system approach that relies on multiple-sequence alignments while ESMFold uses language models. individual plots show the distribution of average residue confidence scores (PLDDT) for each of the six *Fusarium* species. AlphaFold predicted a greater number of proteins with a high confidence score (≥ 70 pLDDT) but takes up to 60 times longer to make each prediction.

### Pan-genome framework with variant effect scores

A pan-genome for 22 diverse Fusarium species (listed in Table 1) was constructed using OrthoFinder [22], and protein sequences downloaded from UniProt [28]. The analysis resulted in 29,529 orthologous clusters (OrthoGroups), which constitute the pan-genome for this set of species and serve as a comprehensive framework for genetic diversity within the genus. Each OrthoGroup was subjected to multiple sequence alignments using FAMSA [29]. Finally, the variant effect scores were calculated using the Evolutionary Scale Modeling (ESM1b) protein language model [26] via the esm-variants tool. The scores predict the functional impact of amino acid substitutions. Scores above −7 predict a benign effect of an amino acid substitution, whereas scores below −7 predict a significant functional change. The variant effect scores were calculated for all 20 possible amino acid substitutions at each position in the two reference proteomes. This analysis encompassed over 127 million potential missense mutations in the *F. graminearum* proteome and over 142 million in the *F. verticillioides* proteome, revealing the vast potential for genetic variation and its significant impact on phenotypic traits. These scores were cross-linked to the naturally occurring amino acid substitutions within the *Fusarium* pan-genome. FPT uses the PanEffect framework [30] to visualize and compare the variation to the reference proteomes, offering insights into the functional consequences of genetic diversity.

Notably, of the 127 million potential missense mutations in *F. graminearum*, fewer than 28% (approximately 35.5 million) are observed within the pan-genome proteins, and for the 142 million possible variants in *F. verticillioides*, only 23% or 32.9 million are found in the pan-genome. See Figure 2 for the distribution of the potential variant effect scores in *F. graminearum* and *F. verticillioides* alongside the distribution of naturally occurring variant effect scores in the *Fusarium* pan-genome.

**Figure 2.**
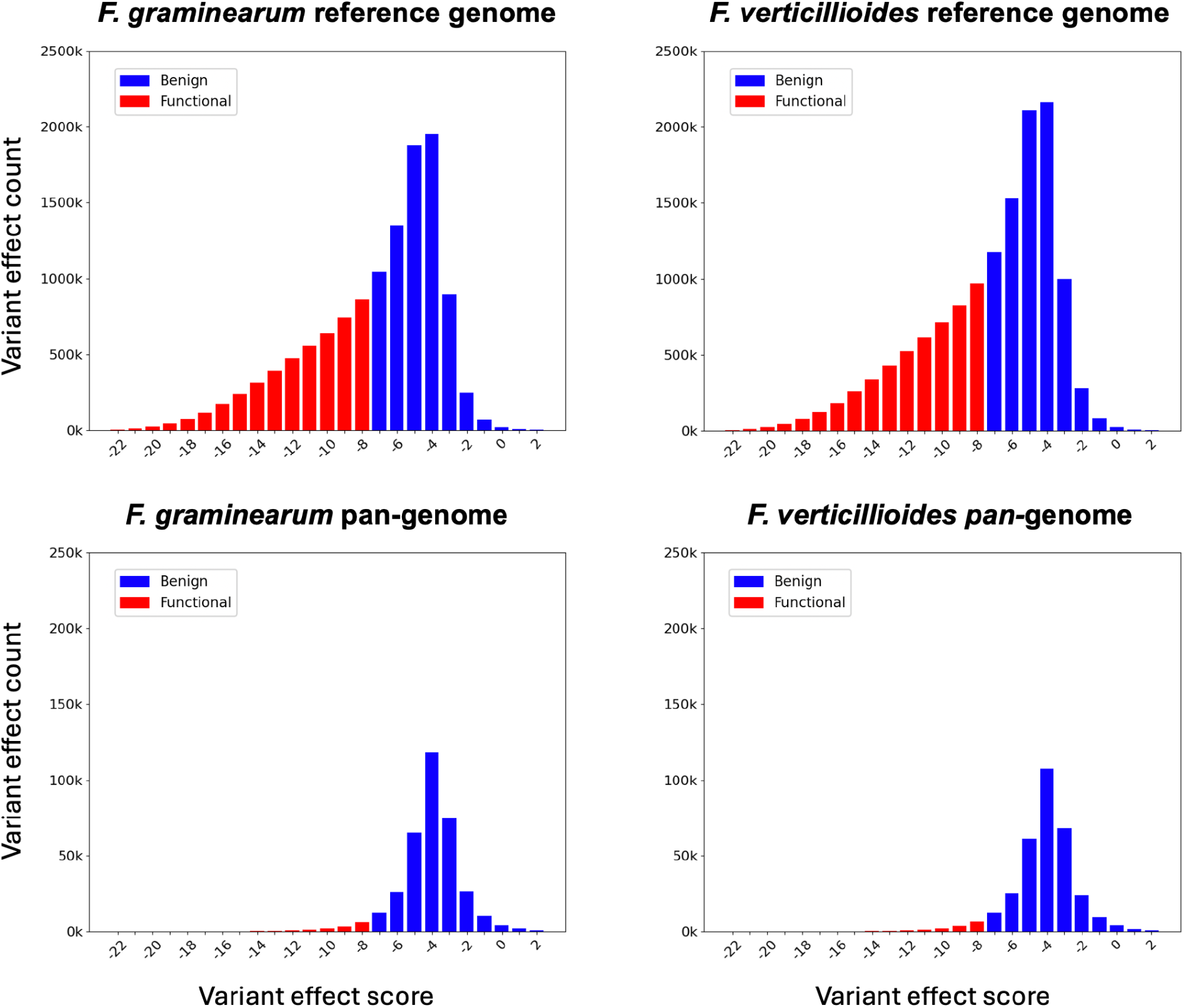
The distribution of variant effect scores in *Fusarium graminearum* and *Fusarium verticillioides*. The upper left panel shows the distribution of the variant effect scores for over 127 million missense mutations among *F. graminearum* proteins. The upper right panel shows the distribution of the variant effect scores for over 142 million possible missense mutations among *F. verticillioides* proteins. The bottom two panels illustrate the distribution of the variant effect scores of the actual missense mutations between the reference proteins and the 22 proteomes in the Fusarium pan-genome. For each panel, the x-axis is labeled by the variant scores and the y-axis shows the count of variants with that score. Red bars have scores less than −7 and are considered likely to have a functional effect. The blue bars have scores greater than or equal to −7 and are more likely to be benign.

## 3 Utility and discussion

The Fusarium Protein Toolkit (FPT) interface (https://fusarium.maizegdb.org/) provides access to a set of tools that enable users to navigate and explore *Fusarium* protein structures and annotations. The underlying datasets are freely available for download through a link provided on the webpage. The toolkit is organized into five sections: Home, Effectors, FoldSeek, PanEffect, and Help, each designed to facilitate specific aspects of *Fusarium* protein research. Table 2 lists the tool and data availability for each of the *Fusarium* species.

### Home page overview

The FPT’s landing page is the home page, which provides quick access to the toolkit’s tools and features. Each section of the home page provides a concise overview of the toolkit’s capabilities with interactive components to search, download, or visualize the underlying data. The Home page incorporates direct search functionalities for the FoldSeek and PanEffect tools, allowing users to quickly query *Fusarium* gene or protein names. The table of annotated predicted effector genes is available through the menu or the “Fusarium Effector webpage” button on the homepage. The download section provides a bulleted list of downloadable datasets. The final two sections provide interactive widgets that display AlphaFold or ESMFold protein structures.

### Effectors page overview

The Effectors webpage provides a table of putative effector proteins for the six representative species of *Fusarium*: *F. graminearum, F. fujikuroi, F. oxysporum, F. proliferatum, F. vanettenii, F. verticillioides*. The tables are organized to show the confidence of the predictions, where the effector proteins are likely to be localized, functional annotations, and direct links to the other tools in the FPT. More specifically, each table includes the protein name, providing a primary reference identifier; linked UniProt IDs for in-depth protein information; apoplastic or intracellular localizations with the prediction probabilities and the amino acid positions of the domains, which are important for understanding effector modes of action; and additional localization details regarding the chloroplast, mitochondria, or nucleus to provide insight to the potential impacts on cellular processes. The table lists functional descriptions that give a concise overview of each protein’s potential functions and roles in *Fusarium*-host interactions. These descriptions are complemented by Gene Ontology (GO) [31] terms that give controlled vocabulary terms for biological processes, cellular components, and molecular functions. When applicable, enzyme codes are listed to indicate enzymatic activities. The final column in the table has direct links to FPT’s PanEffect and FoldSeek tools, alongside links to the AlphaFold Protein Structure Database. These links provide quick access to functional analysis and structural prediction which facilitate a streamlined and integrated workflow for researchers focused on plant pathology, mycology, and plant-microbe interactions. By centralizing curated data of *Fusarium* effector proteins, the Effectors webpage enables users to quickly access detailed information, compare effector functions and localizations, and explore links to external databases for extended research. The integration of direct links to analytical tools and databases provides a transition from gene identification to functional and structural analysis, which is important to advance our understanding of *Fusarium* effector proteins and their roles in plant disease.

### FoldSeek (protein structure search) overview

The FoldSeek Search Tool is designed to provide insights into protein structures and sequence alignments (Figure 3). Utilizing the FoldSeek software, this resource enables the comparison of AlphaFold protein structure alignments from either *F. graminearum* or *F. verticillioides* with nine proteomes. This selection of proteomes includes five *Fusarium* strains and four outgroup proteomes from species such as *A. thaliana* and *H. sapiens*, facilitating a broad comparative analysis. The tool offers visualizations of protein structure alignments, including 3D views of *Fusarium* protein structures predicted by AlphaFold, to help in understanding protein configurations. It also provides access to detailed protein information, such as UniProt IDs, gene names, Pfam domains, and functional annotations. The tool presents structural alignment quality in a color-coded format, making it easier to identify the level of similarities and differences between orthologous proteins. An interactive feature allows users to click on these color-coded bars to access more detailed comparisons of sequences and the superposition of the structural alignments between different species.

**Figure 3.**
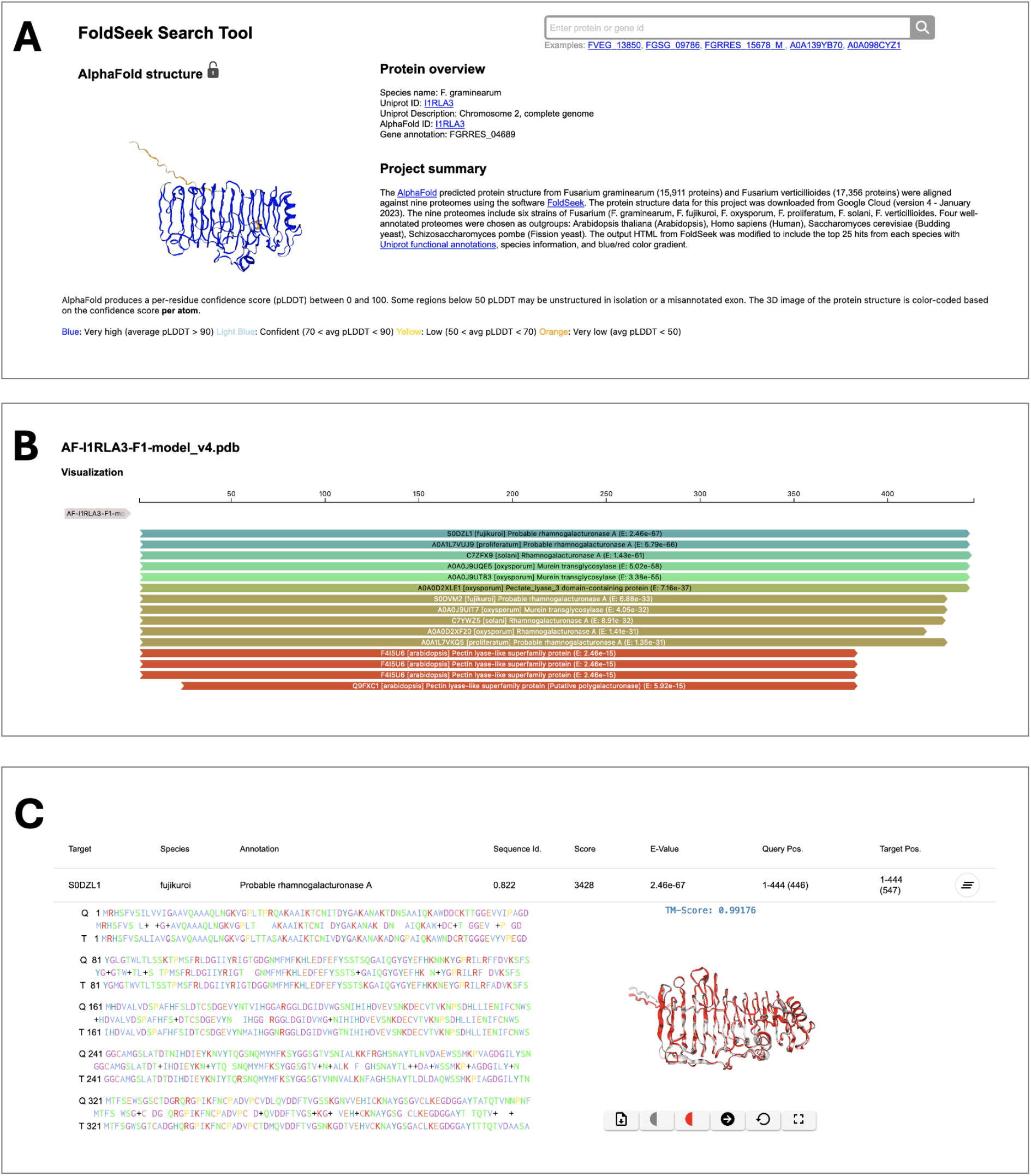
Example page of Fusarium FoldSeek search tool for the *F. graminearum* gene FGRRES_04689 (UniProt: I1RLA3). (A) The protein overview section on the top of the page, which includes the Project summary, Protein overview, and AlphaFold structure. (B) A zoomed-in view of the scores for the 15 top hits of UniProt proteins aligned to FGRRES_04689; the top hit is the S0DZL1 protein from the *F. fujikuroi* annotated as a “Probable rhamnogalacturonase A.” Clicking on the panel containing the alignment score for takes the user to the protein sequence alignment. (C) The structural superposition of the top hit of the *F. graminearum* and *F. fujikuroi* proteins. The table provides additional metrics on the alignment and an interactive structure view.

The FoldSeek Search Tool aids researchers by providing a platform to visualize, analyze, and compare protein structure alignments. This tool will be particularly useful for comparative studies of fungi by identifying orthologs, inferring function, exploring structural domains, and comprehending the evolutionary relationships among proteins that were not evident through sequence alignments alone.

### PanEffect (variant effect viewer) overview

The PanEffect tool (Figure 4) is a resource designed for in-depth analysis of the potential impacts of missense variants on *Fusarium* proteins. It provides users with an interactive interface comprising four specialized views. The Search function enables users to make queries based on gene names and protein identifiers. The Gene Summary section presents an overview of the gene and protein annotations. In the Variant effects within a genome view, detailed heatmaps show the predicted functional impact of all possible amino acid substitutions for *F. verticillioides* or *F. graminearum* proteins. These heatmaps, which transition from blue to red to indicate impact severity, provide insights into each position of the reference protein and the possible amino acid substitutions. Additionally, the tool allows for the exploration of Variant Effects across the pan-genome with additional heatmaps that reflect the effects of natural variations on *Fusarium* proteins across orthology-based gene families built on 22 *Fusarium* species. In addition, over 3 billion effect scores for nearly 330,00 proteins across the 22 species are available as downloads.

**Figure 4.**
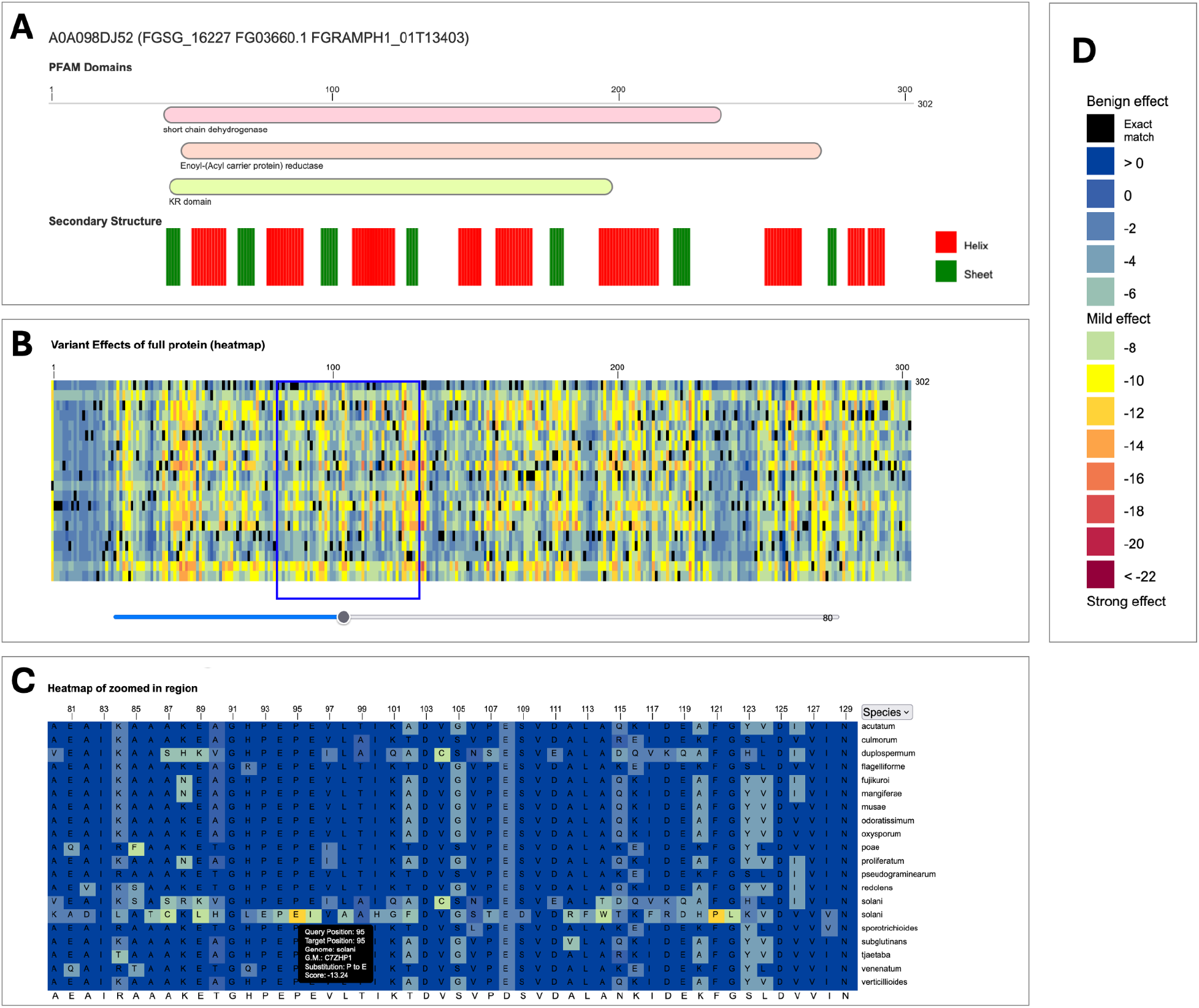
Example page of Fusarium PanEffect tool for the *F. graminearum* gene FGSG_16227 (UniProt: A0A098DJ52). The PanEffect tool has two major views showing the heatmaps of variant effect consequence scores at the reference genome level and pan-genome level. (A) A snapshot of the Pfam domains and predicted secondary structures for the given protein. This view is available on each page to give some functional and structural context to the coding variants. (B) A snapshot of the zoomed-out heatmap view of the “Variants effects within a genome” tab. It displays a heatmap of all possible coding variants color-coded based on how likely it will affect the protein’s function. (C) A snapshot of the zoomed-in heatmap view of the “Variants effects within gene families” tab. It shows a heatmap of the naturally occurring variants found in a gene family across 22 diverse *Fusarium* genomes, where variants that are the same as the reference genome (in this case *F. graminearum)* are shown in dark blue. All other variants are color-coded based on how likely it will affect the protein’s function. Mousing over a position on the heatmap shows additional details including the amino acid substitution and predicted score. (D) The legend of the variant effect scores where scores above −7 indicate benign outcomes, while scores below −7 suggest possible phenotypic effects.

The PanEffect tool leverages protein language models to provide a nuanced understanding of how missense variants affect *Fusarium* proteins. These insights can empower researchers to predict and interpret the functional consequences of genetic variations. Moreover, the tool can support a wide range of comparative studies and help make predictions on how variation affects proteins involved in *Fusarium*-plant interactions.

### Help page overview

The help section offers summaries of all the FPT components and includes detailed descriptions and links to the data sources, tools, downloads, and references used to develop FPT. The Help page also includes a table of *Fusarium* species used to develop this resource with links to NCBI and UniProt.

### Case Study: Using the Fusarium Protein Toolkit to Analyze the FVEG_03351 gene

A key application of FPT is to visualize coding changes and correlate them with both predicted functional consequences and structural differences. Figure 5 presents the analysis of the gene FVEG_03351 (UniProt: W7M0U3) from *F. verticillioides*, utilizing the capabilities of the Fusarium Protein Toolkit. FVEG_03351 is annotated as a cutinase protein [32], which is an inducible extracellular enzyme secreted by microorganisms capable of degrading plant cell walls. This characteristic underscores its potential role in host-pathogen interactions, particularly in agricultural settings where Fusarium species are known pathogens. The toolkit’s Effector Annotation Pipeline categorizes FVEG_03351 as an effector gene (Figure 5A), suggesting its involvement in pathogenicity by facilitating the degradation of plant host tissues, thereby promoting infection. Utilizing the PanEffect tool (Figure 5B), the cutinase domain and its predicted secondary structures within FVEG_03351 are shown. This tool also displays heat maps that are color-coded to represent variant effect consequence scores. In the “Variant Effects in the Gene Family” tab, these scores reveal the impact of missense mutations across gene family members of other Fusarium species. The protein A0A1C3YND2 from *F. graminearum* has a series of variants from position 180 to the end of the protein (shown in the zoomed-in heatmap) that are predicted to strongly affect its function.

**Figure 5.**
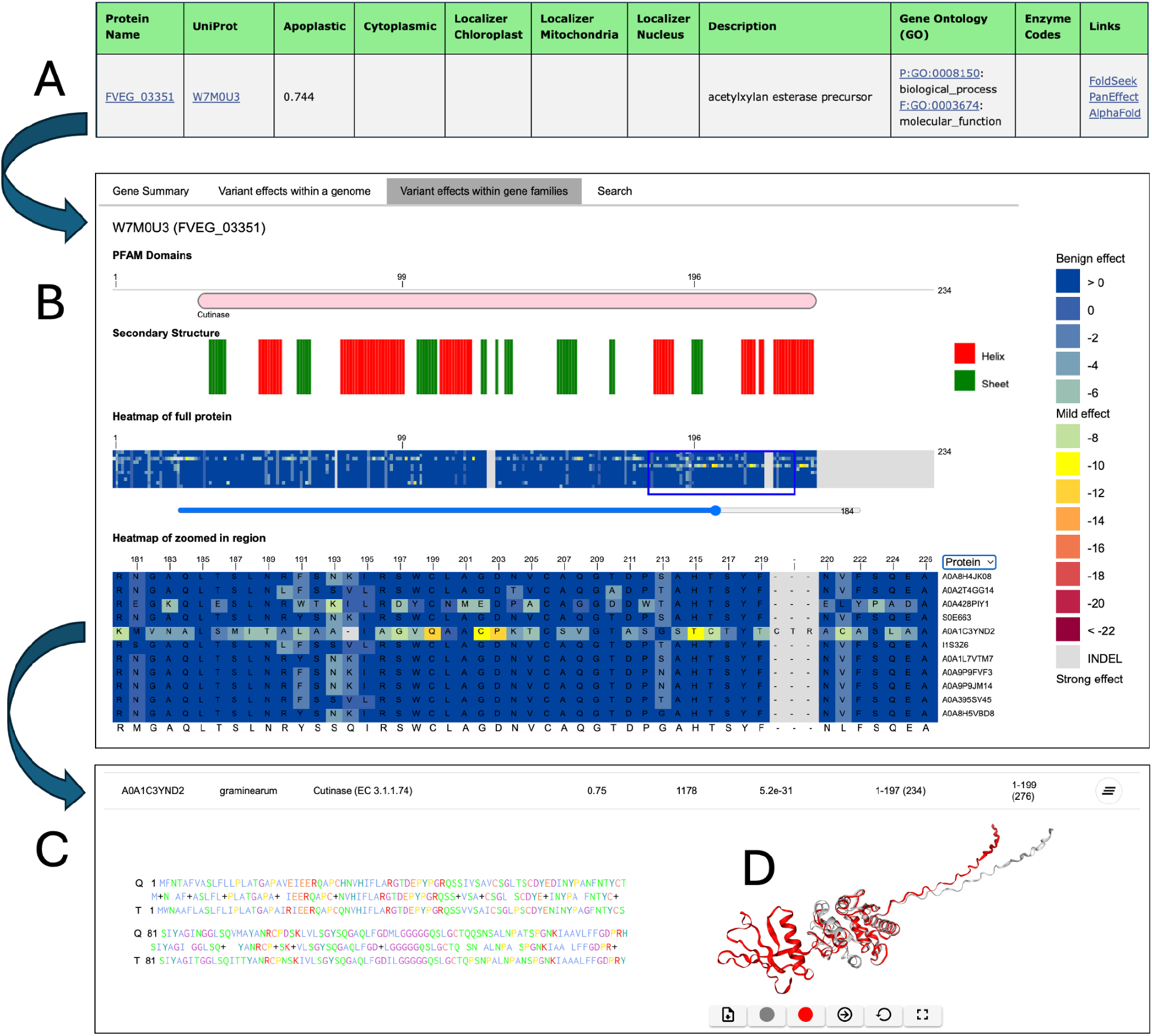
Using the Fusarium Protein Toolkit to Analyze FVEG_03351. The figure shows how the Fusarium Protein Toolkit can be used for the analysis of the FVEG_03351 gene (UniProt: W7M0U3) from *F. verticillioides*. FVEG_03351 is annotated as a cutinase protein, a type of inducible extracellular enzyme secreted by microorganisms that can degrade plant cell walls. A) The Effector annotation pipeline identifies FVEG_03351 as an effector gene. B) The PanEffect tool for FVEG_03351 reveals the presence of the cutinase domain and predicted secondary structures. The tool also displays color-coded variant effect consequence scores in heat maps, including the “Variant effects in the gene family” tab, which shows the consequence scores of missense mutations across 11 members of the gene family that were calculated across 22 *Fusarium* proteomes. In this example, the protein A0A1C3YND2 from *F. graminearum* contains several variants predicted to have a strong functional effect (zoomed-in region). C) A sample output of the FoldSeek tool for proteins W7M0U3 (query protein) and A0A1C3YND2 (target protein) which is also accessible through the Links column in the Effectors table. D). In the structure superposition view, the query protein W7M0U3 is shown in gray, and A0A1C3YND2 in red. The structure shows the portion of A0A1C3YND2 from position 190 to the terminal of the protein conforms to a different structure as W7M0U3.

The FoldSeek tool (Figure 5C) can be used to examine the structural alignments and identify structural changes caused by the coding differences between the query protein W7M0U3 in *F. verticillioides* and the target protein A0A1C3YND2 in *F. graminearum*. This analysis is valuable for understanding the conformational variations that might influence protein function and interaction dynamics. The structural superposition view in the FoldSeek tool (Figure 5D) is depicted with the query protein W7M0U3 in gray and A0A1C3YND2 in red. This comparison visually emphasizes the structural differences between the two proteins, specifically the stretch from position 180 to the terminal end of A0A1C3YND2 which overlaps the region found in PanEffect. The structural differences observed are substantial for this region and provide further support for the possibility of functional or phenotypic differences between the two proteins. This analysis used the FPT to identify structural and functional differences of an important gene implicated in plant disease. By detailing the contributions of each component of the toolkit, this figure underscores the multifaceted approach required to understand the roles of such proteins in pathogenicity, offering insights that are crucial for developing targeted strategies against *Fusarium*-related plant diseases.

## 4 Conclusions

The Fusarium Protein Toolkit should be an important resource in fungal pathogen research by offering a comprehensive suite of tools to explore protein structures, variant effects, and annotated effector proteins of *Fusarium*. Researchers can visualize and analyze protein structures from AlphaFold and ESMFold models, comparing them across diverse Fusarium species and outgroups. The toolkit also leverages protein language models to predict the functional impact of missense mutations, providing valuable insights into how genetic variations might affect protein function. Additionally, the PanEffect tool allows researchers to explore the effects of natural variations within a pan-genome based on 22 *Fusarium* species, aiding in understanding the potential impact of genetic diversity on the biology of these fungi.

By offering these capabilities, FPT will significantly deepen our understanding of the pathogenic mechanisms of *Fusarium*. This information and knowledge have the potential to facilitate the identification of proteins that can be used as targets to control *Fusarium*-incited crop diseases and mycotoxin contamination and thereby ensure food security in a changing climate. FPT takes a step forward not only by offering these *Fusarium*-specific insights but also by being built upon the same frameworks and tools used for the model host species, maize. This shared infrastructure lays the groundwork for future work aimed at understanding host-pathogen interactions. By enabling direct comparisons of protein structures and functionalities between *Fusarium* and maize, researchers can gain a deeper understanding of how these organisms interact at the molecular level. This knowledge can inform the development of more precise strategies to target the vulnerabilities within these interactions, ultimately leading to improved disease management and crop protection.

## Data Availability

Fusarium Protein Toolkit is freely accessible at https://fusarium.maizegdb.org/ and is maintained by MaizeGDB. The PanEffect framework is available at https://github.com/Maize-Genetics-and-Genomics-Database/PanEffect. The underlying data generated from the artificial intelligence and bioinformatics approaches are found in the Fusarium data folder in the Artificial Intelligence section of the MaizeGDB download page (https://maizegdb.org/download).

## Acknowledgements

The work for this project was performed on the Atlas and Ceres high-performance clusters as part of the USDA-ARS SCINet initiative. We would like to thank the SCINet administrative staff and the Virtual Research Support Core team. This work was supported by the U.S. Department of Agriculture, Agricultural Research Service. Mention of trade names or commercial products in this publication is solely for the purpose of providing specific information and does not imply recommendation or endorsement by the U.S. Department of Agriculture. USDA is an equal opportunity provider and employer.

